# Regeneration in the adult *Drosophila* brain

**DOI:** 10.1101/2020.01.16.908640

**Authors:** Kassi L. Crocker, Khailee Marischuk, Stacey A. Rimkus, Hong Zhou, Jerry C.P. Yin, Grace Boekhoff-Falk

## Abstract

Neurodegenerative diseases such as Alzheimer’s and Parkinson’s currently affect ∼25 million people worldwide (Erkkinen *et al*. 2018). The global incidence of traumatic brain injury (TBI) is estimated at ∼70 million/year (Dewan *et al*. 2018). Both neurodegenerative diseases and TBI remain without effective treatments. We are utilizing adult *Drosophila melanogaster* to investigate the mechanisms of brain regeneration with the long term goal of identifying targets for neural regenerative therapies. Like mammals, Drosophila have few proliferating cells in the adult brain. Nonetheless, within 24 hours of a Penetrating Traumatic Brain Injury (PTBI) to the central brain, there is a significant increase in the number of proliferating cells. We subsequently detect both new glia and new neurons and the formation of new axon tracts that target appropriate brain regions. Glial cells divide rapidly upon injury to give rise to new glial cells. Other cells near the injury site upregulate neural progenitor genes including *asense* and *deadpan* and later give rise to the new neurons. Locomotor abnormalities observed after PTBI are reversed within two weeks of injury, supporting the idea that there is functional recovery. Together, these data indicate that adult *Drosophila* brains are capable of neuronal repair. We anticipate that this paradigm will facilitate the dissection of the mechanisms of neural regeneration and that these processes will be relevant to human brain repair.

## Introduction

Although classical studies reported that the mammalian brain stops making new neurons shortly after birth (Ramon y Cajal 1913; Ramon y Cajal 1914), populations of dividing progenitor cells subsequently were observed in two major regions of the rodent brain: the subventricular zone (SVZ) of the forebrain and the dentate gyrus (DG) of the hippocampus (Altman and Das 1965; Altman 1969; Kaplan and Hinds 1977). Both SVZ and DG progenitors give rise to multiple cell types over the course of an animal’s lifetime, including neurons, astrocytes and oligodendrocytes (Kuhn *et al*. 1996). Moreover, a landmark study published in 1998 concluded that adult humans create new neurons in the hippocampus in the absence of injury (Eriksson *et al*. 1998).

The discovery of adult neurogenesis raised the possibility of utilizing endogenous cells for neural regeneration both following brain injury and also in patients suffering from neurodegenerative diseases. Although emphasis in the mammalian neural regeneration field to date has been on transplanting embryonic or induced pluripotent stem cells (Vishwakarma *et al*. 2014), obtaining functional integration of transplanted neural cells remains a major challenge. Moreover, stem cell transplants can be accompanied by tumor formation (Amariglio *et al*. 2009). A more recent direction for the field therefore has been on coaxing resident cells in the brain to undertake regeneration (Gao *et al*. 2016).

In order to utilize endogenous cells for neural repair in clinical settings, we first must identify the cell types and underlying molecular mechanisms that contribute to neuroregeneration. To address these questions, we are investigating repair of the adult brain in the fruitfly *Drosophila melanogaster.* The adult *Drosophila* brain, like the adult mammalian brain, has few proliferative cells (von Trotha *et al*. 2009). In addition, although the *Drosophila* brain has many fewer neurons, the *Drosophila* brain has many shared complexities with the human brain including analogous neural cell types (Lessing and Bonini 2009), common neurotransmitters (GABA, glutamate, and acetylcholine) (Bellen *et al*. 2010), similar synapse architecture (Lessing and Bonini 2009) and similar physiology and intracellular signaling pathways (Bellen *et al*. 2010). These properties, combined with a wealth of genetic and molecular tools, a short generation time, and large number of offspring, lead us to propose that *Drosophila* offer an exceptional model in which to investigate brain regeneration.

The combination of rare cell proliferation and the fact that known neural progenitors undergo terminal differentiation or apoptosis during metamorphosis (Siegrist *et al*. 2010), make it all the more remarkable that the adult *Drosophila* brain is capable of neurogenesis. Nonetheless, the adult *Drosophila* brain can make new neurons after injury (Fernandez-Hernandez *et al*. 2013; Moreno *et al*. 2015). These earlier studies focused on the adult optic lobes, where slowly cycling neural progenitor cells were discovered in the medulla cortex. These progenitors are activated by injury and give rise to new neurons (Fernandez-Hernandez *et al*. 2013; Moreno *et al*. 2015).

Here, in contrast to other studies, we focus on a distinct brain region, the central brain. We find that the central brain, like the optic lobes, can produce new neurons after injury. However, in contrast to the optic lobes, the new central brain neurons innervate specific targets, including the mushroom body, ellipsoid body, antennal lobes and lateral horn. We present evidence that these new neurons contribute to functional regeneration. Also, in contrast to the optic lobes, the central brain produces new glial cells in response to injury. We have lineage-traced both the new neurons and the new glia and find that they arise from distinct progenitors. We propose that an injury-triggered mechanism results in the reactivation of neural progenitor genes, allowing these cells to proliferate and give rise to new neurons. Because of the extensive parallels between adult Drosophila and mammalian brains, we anticipate that these studies will have relevance to human neural regeneration.

## Results

### Penetrating Traumatic Brain Injury (PTBI) stimulates cell proliferation

To investigate the regenerative capacity of the adult *Drosophila melanogaster* central brain, we examined brains at a variety of time points following a penetrating traumatic brain injury (PTBI). We injured the central brain near the mushroom body (MB; **Fig. 1A**) through the head cuticle using a stainless steel insect pin (∼12.5μm diameter tip, 100 μm diameter rod). This is the same type of pin used by Fernandez-Hernandez *et al*. to injure adult brains (Fernandez-Hernandez *et al*. 2013), but distinct from that used by Sanuki *et al*. (Sanuki *et al*. 2019) who used larger (150 μm diameter), hollow, syringe needle. These earlier studies examined the consequences of injury to the adult optic lobes. In contrast, we injured the adult central brain near the mushroom body. Mushroom body neuroblasts are the last to stop proliferating during development (Ito and Hotta 1992; Ito *et al*. 1997; Siegrist *et al*. 2010). We therefore reasoned that the mushroom body might have residual mitotic potential. Located dorsally in the central brain, the mushroom body is critical for learning and memory and contains complex dendrite and axon arbors in highly stereotyped arrays (Aso *et al*. 2014).

**Figure 1.**
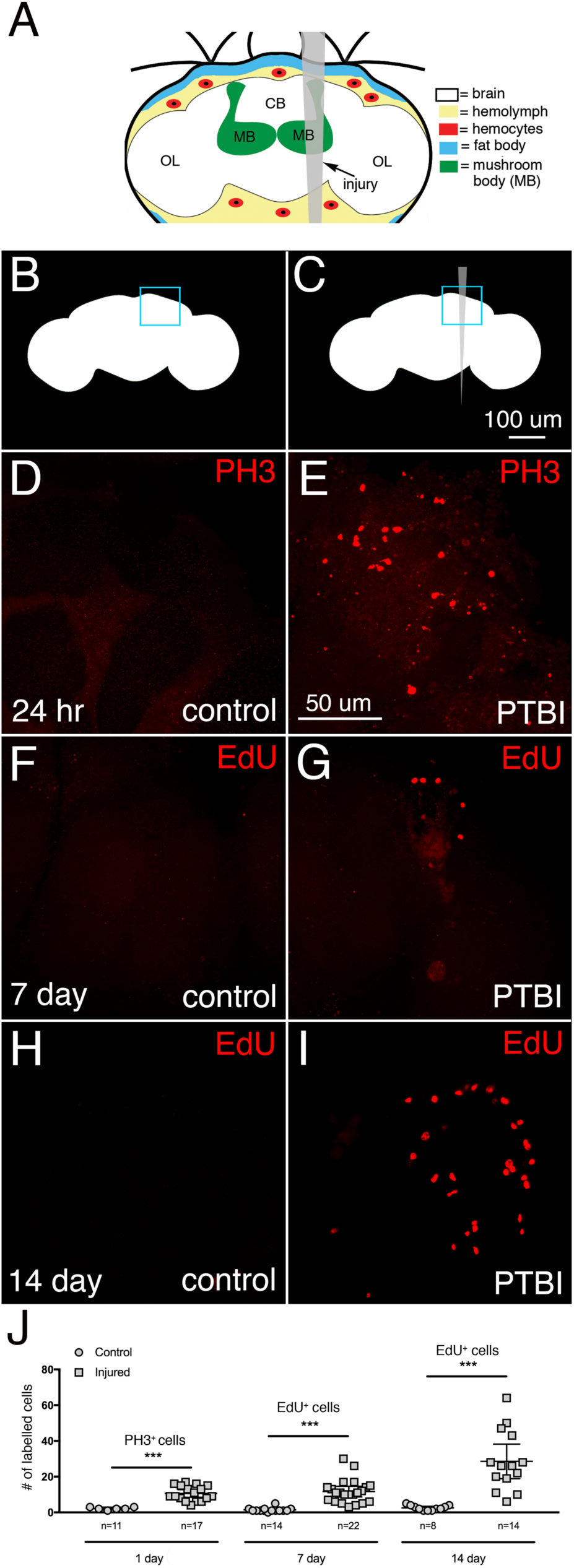
Penetrating Traumatic Brain Injury (PTBI) stimulates cell proliferation. **A.** Schematic of an adult Drosophila head with the injury trajectory indicated in grey. Central brain PTBI impacts multiple brain structures including the mushroom body (MB, green), and tissues outside the brain including the fat body (blue) and hemocytes (red). CB = central brain region. OL= optic lobe region. Uninjured, control (**B**) and PTBI **(C)** schematics. The blue boxes in the upper right corners indicate the brain regions shown at higher magnification in panels **D-I. D**, **E**. PH3 antibody (red) was used to assay for cell proliferation 24 hours after injury. In control brains (**D)** there are few PH3+ cells, and none near the MB. However, in PTBI brains (**E**), there are PH3+ cells near the MB. **F, G.** To test whether newly created cells are surviving or being eliminated, we conducted a pulse-chase EdU experiment. Flies were fed EdU (red) for 4 days post-injury (a pulse) then chased for 3 days without feeding EdU. In the control brain (**F**), there is little EdU incorporation. In the PTBI brain (**G**), there are EdU+ cells near the MB. **H.** In 14-day control brains, there are few EdU+ cells. **I.** However, in 14-day PTBI brains, there is an increase in EdU+ cells near the MB. All brains are from males of our standard genotype. **J.** Quantification of proliferating cells. At 24 hours, uninjured control brains had an average of 3 PH3+ cells/brain (n=11 brains, 28 cells), while 24-hour post-PTBI brains had an average of 11 PH3+ cells/brain (n=17 brains, 181 cells). At 7 days, uninjured controls have few EdU+ cells, with an average of 2 EdU+ cells/brain (n=15 brains, 24 cells), while 7-day post-PTBI brains had an average of 11 EdU+ cells/brain (n=22 brains, 238 cells). At 14 days, uninjured controls have an average of 1 EdU+ cell/brain (n=8 brains,11 cells), while 14-day post-PTBI brains have an average of 29 EdU+ cells/brain (n=14 brains, 400 cells). Unpaired t tests of control and PTBI samples at the 3 time points yield values of p<0.0001, p<0.0001 and p<0.0002, respectively. Error bars reflect the standard deviation (SD).

One early hallmark of a regenerative response is cell proliferation (Zhou *et al*. 2016). We therefore tested whether PTBI stimulates proliferation, using two different assays. We first utilized antibodies to the mitotic marker, phospho-histone H3 (PH3). Histone H3 is transiently phosphorylated during M phase of the cell cycle (Hans and Dimitrov 2001), and antibodies to PH3 are often used to identify dividing cells. Within 24 hours of PTBI, brains from young adult males exhibited a significant increase of PH3+ cells compared to uninjured controls of the same sex, age, and genotype (**Fig. 1B-E, J**). We then wanted to determine whether these newly created cells were maintained or eliminated. Because PH3 only transiently labels dividing cells, we also used incorporation of the nucleotide analog 5-ethynyl-2’-deoxyuridine (EdU). EdU is incorporated into newly synthesized DNA and therefore more permanently labels dividing cells and their progeny. After feeding flies EdU for various lengths of time, we detected the incorporated EdU using fluorescent ‘click chemistry’ (InVitrogen®). Consistent with the PH3 labeling at 24 hours, we observed 5- and 15-fold more EdU+ cells at 7 days and 14 days post-injury, respectively, compared to uninjured controls (p-value<0.0001 and p-value<0.0002) (**Fig. 1F-J**). This indicates that cell proliferation continues between 7 and 14 days post-PTBI.

EdU permanently marks dividing cells and allows us to measure the cumulative number of cells that had divided since the PTBI. In contrast, PH3 provides a snapshot of actively dividing cells. To evaluate the dynamics of cell proliferation, we therefore compared mitotic activity using anti-PH3 in animals at 24 hours, 7 days and 14 days post-PTBI (**Fig. S1A, E**). Consistent with the EdU labeling experiments above, we observed actively dividing cells even 14 days post-injury, although the numbers of dividing cells decrease with time post-PTBI. To test whether age impacts regenerative capacity, we aged adult males to 7, 14, and 28 days post-eclosion prior to PTBI, then assayed for cell proliferation 24 hours later.

Adult male brains injured 7 days post-eclosion had significantly fewer PH3+ cells than brains injured within 6 hours of eclosion (**Fig. S1B**). The brains injured at 7 days had more PH3+ cells than the age-matched uninjured controls, but fewer PH3+ cells than brains injured within 6 hours of eclosion (**Fig. S1B**). Brains injured at either 14 or 28 days post-eclosion exhibited little cell proliferation that did not differ from age-matched uninjured controls (**Fig. S1B**). Taken together, these data support the idea that early in adulthood, *Drosophila* brains possess cells that can initiate division in response to injury and that this proliferative ability declines with age.

The increase in EdU+ cells between 7 and 14 days post-PTBI (**Fig. 1J**) and the presence of actively dividing, PH3+ cells at 7 and 14 days post-PTBI (**Fig. S1A**) indicate that the capacity for sustained cell proliferation is retained beyond the time window during which cell proliferation can be initiated. In other words, once adult brain cells begin to divide, they may continue to divide for up to 2 weeks. However, by 2 weeks of adulthood, brain cells no longer can be stimulated by injury to divide. Of note, uninjured flies at both 24 hours and 7 days exhibit similar baseline cell proliferation (**Fig. S1B**). Taken together, this indicates that the presence of PH3+ cells in uninjured brains does not correlate directly with the brain’s regenerative potential.

Pulse-chase experiments allowed us to test whether the cells incorporating EdU were viable; if a cell dies after synthesizing DNA, the incorporated EdU is expected to be either diffuse or punctate instead of uniformly distributed within each labeled nucleus. There were similar numbers of labeled nuclei in brains from animals either pulse-chase or continuously fed EdU (**Fig. S1C**). This indicates that most EdU-labeled cells survive. Interestingly, at later timepoints post-PTBI, there was a statistically significant increase in the number of EdU-positive cells not only in the injured brain hemisphere, but also in the contralateral, uninjured brain hemisphere (**Fig. S1D**). This indicates that PTBI may induce both local and widespread proliferation. For this reason, we use brains from uninjured animals as controls instead of the contralateral, uninjured brain hemispheres.

### Characterizing the identities of dividing cells post-injury

To determine the identities of the mitotically active cells, we fed young adult males with EdU, then simultaneously assayed for EdU incorporation, expression of the glial protein Reversed polarity (Repo) and expression of the neuronal protein Embryonic lethal, abnormal vision (Elav). At 7 days post-injury, we observed 4 classes of cells (**Fig. 2A-D’’’**): cells that were EdU+ but did not express either Elav or Repo (arrowheads in **Fig. 2A-A’’’**; Class I); cells that were EdU+ and Repo+, i.e. glia (arrowheads in **Fig. 2B-B’’’**; Class II); cells that were EdU+ and express Elav, i.e. neurons (arrowheads in **Fig. 2C-C’’’**; Class III); and cells that were EdU+, Repo+, and Elav+ (arrowheads in **Fig. 2D-D’’’**; Class IV). These data indicate that cells actively divide after PTBI, and that the dividing cells either are, or become, glia and neurons.

**Figure 2.**
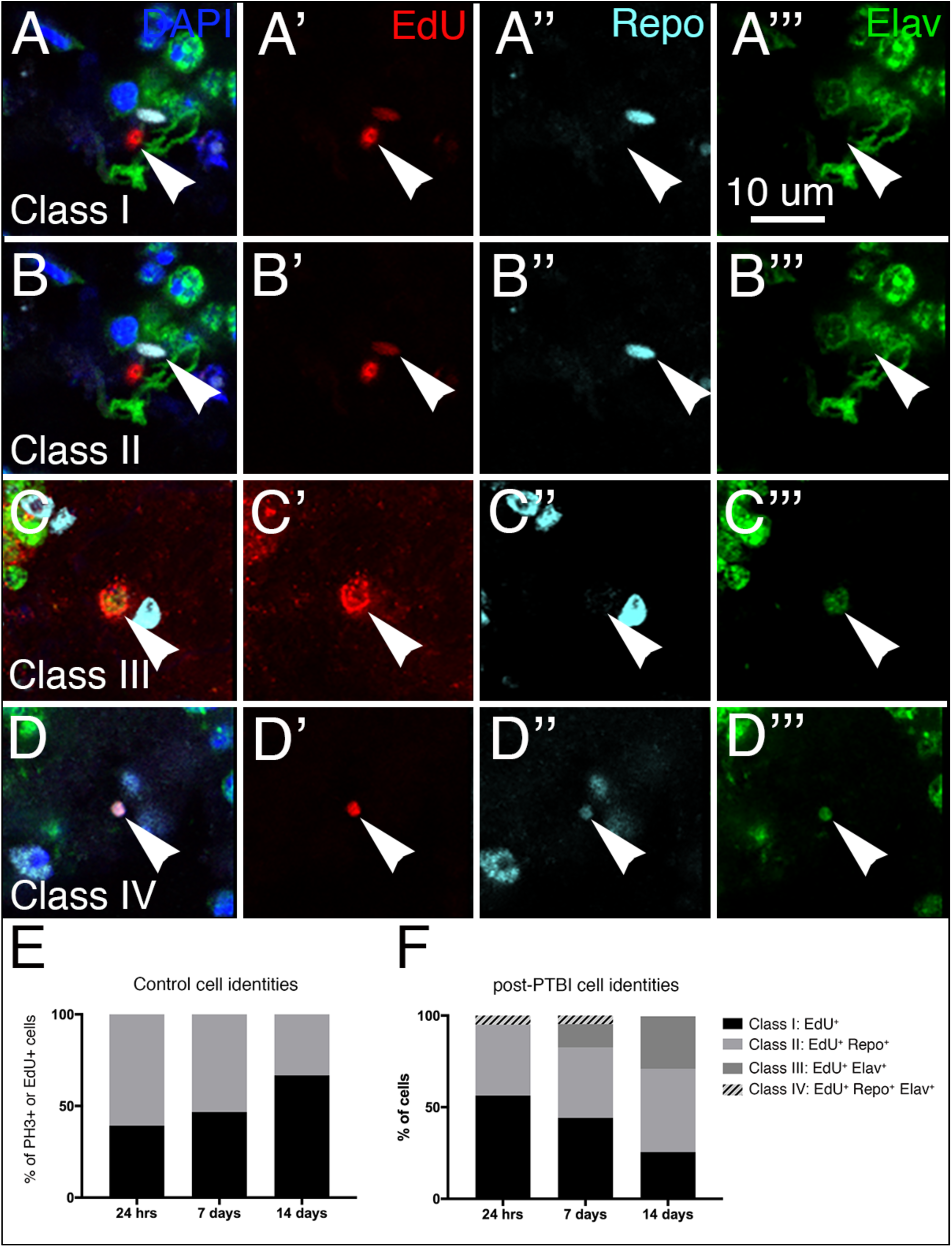
Analysis of new cell identities 7 days post-PTBI. To determine what types of new cells are generated in the first 7 days post-PTBI, we used pulse-chase experiments with EdU in combination with the glial marker anti-Repo and the neuronal marker anti-Elav. At 7 days post-PTBI, we found four classes of cells resulting from proliferation: EdU+ and without either Repo or Elav (**A-A’’’**; Class I); Cells that were EdU+ and Repo+ (**B-B’’’**; Class II); EdU+ and Elav+ (**C-C’’’**; Class III); and EdU+, Repo+, Elav+ (**D-D’’’**; Class IV). Arrowheads indicate representative cells in each class. The nuclear dye DAPI is in blue. We then measured the changing ratios of these cell types over time in control (**E**) and PTBI brains (**F**). For controls, 11 brains and 28 PH3+ cells were analyzed at 24 hours, 9 brains and 15 EdU+ cells were analyzed at 7 days, and 2 brains and 3 EdU+ cells were analyzed at 14 days. For PTBI samples, 17 brains and 181 PH3+ cells were analyzed at 24 hours, 15 brains and 172 EdU+ cells were analyzed at 7 days, and 8 brains and 278 EdU+ cells were analyzed at 14 days.

As described above, age plays an important role in the proliferative capacity of brain cells. We therefore asked whether the amount of time after injury impacts the identities of proliferating cells. Specifically, we compared the ratios of proliferating cells in each of the 4 classes at 24 hours, 7 days and 14 days post-PTBI. (**Fig. 2F**). The proliferation marker we used in the 24-hour assay was anti-PH3, while the marker used in the 7 and 14 day assays was EdU incorporation. At 24 hours post-injury, we observed the following distribution of classes: 56% Class I (PH3+); 38% Class II (PH3+Repo+); and 5% Class IV (PH3+Repo+Elav+) (**Fig. 2F**). Significantly, at 24 hours post-PTBI, there are no Class III (PH3+Elav+) cells (**Fig. 2F**). By 7 days post-PTBI, 13% of the EdU+ cells are Class III, i.e. new neurons (**Fig. 2F**). By 14 days post-PTBI, 29% of the EdU+ cells are new neurons (**Fig. 2F**). Also, by 14 days post-PTBI, there are no longer any Class IV cells that express a hybrid glial and neuronal identity. Of significance, neither Class IV cells, nor the Class III adult-generated neurons are detected in uninjured adult brains at any of the three timepoints (**Fig. 2E)**. Also, although the ratios of Class I to Class II cells are similar in uninjured and injured brains, the total numbers of dividing cells are quite different. 24 hour control brains averaged 2.5 dividing cells/brain (11 brains; 28 PH3+ cells) compared to 10.7 dividing cells/brain following PTBI (17 brains; 181 PH3+ cells). At 7 days, control brains averaged 1.7 EdU+ cell/brain (9 brains; 15 dividing cells) compared to 11.5 EdU+ cells/brain following PTBI (15 brains; 172 EdU+ cells). And at 14 days, control brains averaged 1.5 EdU+ cells/brain (2 brains; 3 dividing cells) compared to 34.8 EdU+ cells/brain following PTBI (8 brains; 278 EdU+ cells). Together these data demonstrate: 1) that glia can divide to give rise to new glia; 2) that new neurons are created later than new glial cells; and 3) that some of the Class I (EdU+) and/or Class IV (EdU+Repo+Elav+) cells may give rise to new neurons (EdU+, Elav+). For reasons explained below, we favor the possibility that it is the Class I cells that give rise to the new neurons.

### Expression of neural progenitor genes post-PTBI

Although neural stem cells have been reported in the optic lobes of the adult Drosophila brain (Fernandez-Hernandez *et al*. 2013), there are no known neural stem cells (neuroblasts) in the adult Drosophila central brain. To test whether neuroblast fates were induced by injury and, if so, whether the generation of new neurons followed a normal developmental trajectory, we assayed the expression of multiple neural precursor genes post-PTBI using immunohistochemistry and/or qRT-PCR. During normal Drosophila development, the central brain neurons and glia derive from two types of neuroblasts (Boone and Doe 2008; Egger *et al*. 2008; Weng and Lee 2011; Homem and Knoblich 2012; Homem *et al*. 2015)2008. Type I neuroblasts express the transcription factors Deadpan (Dpn) and Asense (Ase). Type II neuroblasts express Dpn, but not Ase, and give rise to intermediate neural progenitors (INPs) (Bello *et al*. 2008; Boone and Doe 2008; Bowman *et al*. 2008). Immature INPs express the transcription factor Ase, but not Dpn while mature INPs no longer express Ase, but reactivate Dpn. Thus there is coexpression of Dpn and Ase in Type I, but not in Type II lineages. Other transcription factors required during neurogenesis in the Type I and/or Type II lineages include Inscuteable (Insc) and Earmuff (Erm) (Chia *et al*. 2008).

In immunohistochemistry experiments, we probed control and injured brains from *ase-Gal4, UAS-Stinger; UAS-Gal80^ts^* adult males with anti-Dpn and anti-PH3 antibodies. These flies were reared at 18°C where the temperature-sensitive Gal80 protein is functional and prevents expression of green fluorescent protein (GFP) from the Stinger construct (Barolo *et al*. 2000). Within 6 hours of eclosion, adult male flies were subjected to PTBI and then placed at 30°C for 24 hours prior to dissection and immunostaining. At 30°C, the temperature-sensitive Gal80 is not functional and GFP is expressed in *ase*-expressing cells. We observed clusters GFP+ cells and cells that are PH3+Dpn+ in PTBI, but not control, brains (**Fig. 3A-D’’’**). This indicates that proliferating cells have a key feature of neuroblast identity, namely Dpn expression. We did not observe cells that were PH3+Dpn+GFP+, i.e. dividing and coexpressing *dpn* and *ase*. However, the juxtaposition of the PH3+Dpn+ cells to cells expressing *ase* is reminiscent of Type II neuroblast lineages and consistent with the PH3+Dpn+ cells and Ase+ cells sharing a common origin.

**Figure 3.**
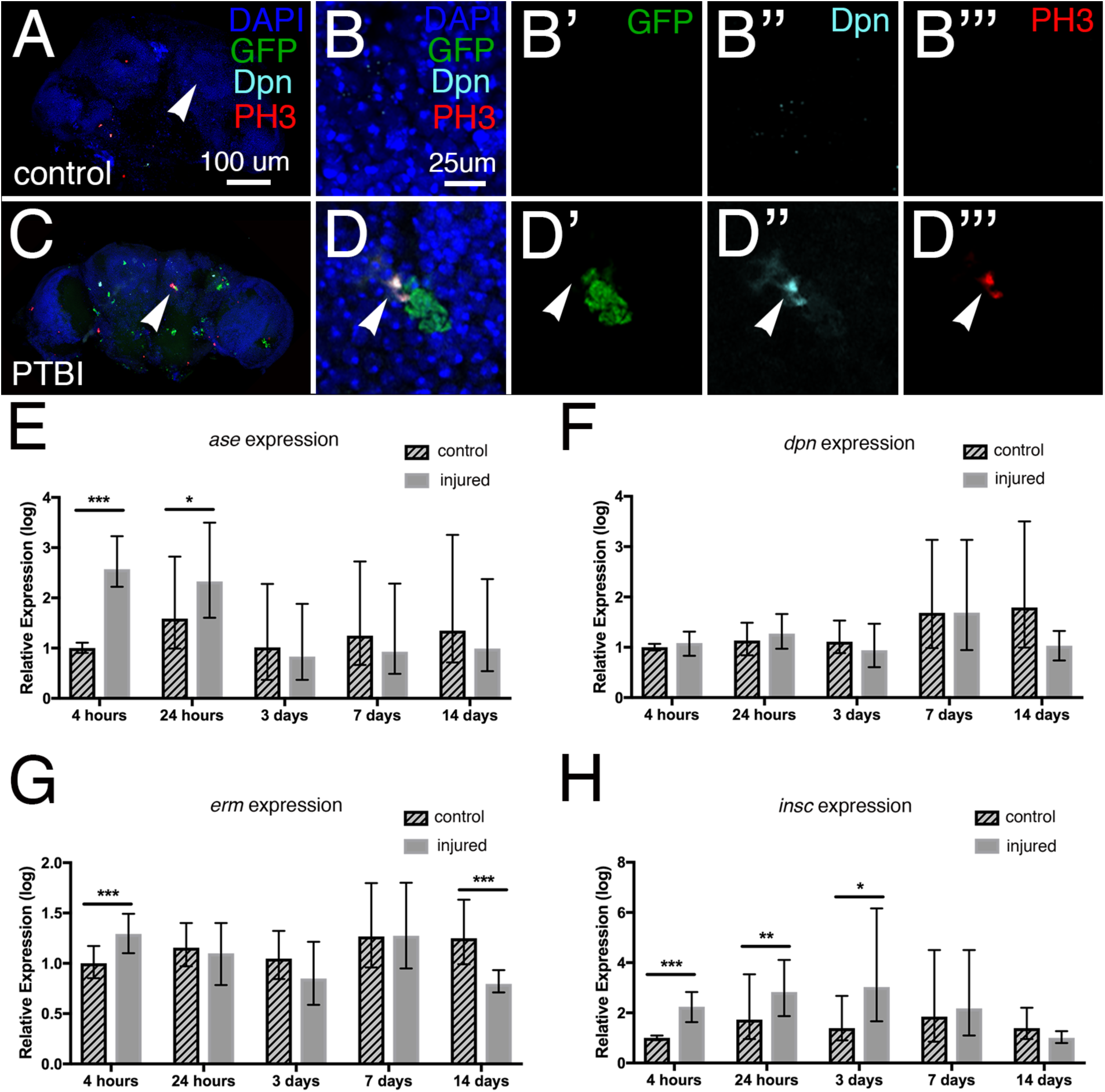
Neuroblast gene expression is activated by PTBI. (**A**) and (**C**) are low magnification views of the control and PTBI brains shown at higher resolution in **B-B’’’** and **D-D’’’**, respectively. **A-B’’’.** Images from an uninjured *ase-Gal4, UAS-Stinger; UAS-Gal80^ts^* probed with anti-PH3 (red) and anti-Dpn (cyan). GFP (green) from the *UAS-Stinger* construct is expressed under control of *ase* regulatory sequences. The nuclear dye DAPI is in blue. Arrowheads in (**A**) and (**C**) indicate the regions where higher magnification images were collected. Animals were reared at 18°C where the temperature-sensitive Gal80 repressor is active and shifted to 29°C after eclosion to permit expression of GFP in cells expressing *ase*. At 24 hours post-PTBI (**C-D’’’**), but not in control (**A-B’’’**) brains, there are GFP+ cells, indicating the expression of *ase*, which is a neural progenitor gene. (**D-D’**). Cells that were Dpn+ (cyan) and PH3+ (red) also were observed in injured brains (**D’’-D’’’**), but not in controls (**B’’-B’’’**). *dpn* is a neuroblast and neural progenitor gene. The PH3+Dpn+ cells were often in close proximity to GFP+ cells (**D**), consistent with a lineal relationship.**E-H.** qRT-PCR reveals increases in neural progenitor gene expression following PTBI. The mRNA levels of four different neural progenitor genes were assayed at 4 hours, 24 hours, 3 days, 7 days, and 14 days. **E.** The level of *ase* mRNA is increased more than 5-fold by 4 hours and remains elevated at 24 hours. However, at 3, 7, and 14 days, *ase* mRNA levels are no longer higher than in controls. **F.** The level of *dpn* mRNA was not detectably increased at any timepoints. **G.** mRNA levels of *erm* are increased almost 3-fold at 4 hours post-injury. However, by 24 hours, 3 days, and 7 days, *erm* mRNA levels have returned to baseline. **H.** *insc* mRNA levels are increased 6-fold at 4 hours, 24 hours, and 3 days post-injury. At later timepoints, 7 and 14 days, *insc* mRNA levels return to near baseline. The qRT-PCR results reflect triplicate biological samples, represented relative to the levels of Rp49, and then normalized to the corresponding levels in time-matched controls. Error bars calculated by Relative Expression Software Tool analysis and reflect the standard error of the mean (SEM). Note that scales on Y axes differ among the graphs.

To quantify neural progenitor gene expression, we collected and injured young adult males within 6 hours of eclosion. mRNA subsequently was extracted at 5 different time points (4 hours, 24 hours, 3 days, 7 days, and 14 days) from post-PTBI and age-matched, uninjured control heads. Relative transcript levels of neural progenitor genes were measured using quantitative real-time PCR (qRT-PCR). At 4 and 24 hours of age, *ase* expression is increased greater than 2-fold in injured flies compared to controls (**Fig. 3E**). This difference diminishes by 3 days and is not significant at later timepoints (**Fig. 3E**). Although we observed Dpn+ cells using immunohistochemistry near the area of injury at 24 hours post-PTBI, *dpn* mRNA levels were not detectably increased at any timepoints (**Fig. 3F**). This could be because we isolated RNA from whole heads and relatively few cells activate *dpn* following PTBI making it difficult to detect an increase above baseline. Alternatively, the increase in Dpn protein levels could be due to post-transcriptional events and not correspond to an increase in steady state mRNA levels. Consistent with the idea that neural progenitor-like cell lineages are generated following PTBI, *erm* transcript levels were increased at 4 hours and *insc* transcript levels were significantly increased at 4 hours, 24 hours, and 3 days, while (**Fig. 3G, H**). *erm* is expressed in Type II neuroblast lineages where it prevents reversion of intermediate neural progenitors to neuroblast fates (Weng *et al*. 2010). *insc* is expressed in Type I neuroblast lineages where it prevents their transformation into intermediate neural progenitors of the Type II lineage (An *et al*. 2017). Together these data support the hypothesis that there are neuroblast-like precursor cells in the adult brain that are either induced or activated by PTBI.

### Lineage-tracing to identify the origins of newly created cells

The presence of neuroblast-like cells post-PTBI could result from differentiated cells such as glia or neurons dedifferentiating and adopting neuroblast-like fates; or via activation of a quiescent population of adult neural stem cells, similar to what was described in the optic lobe (Fernandez-Hernandez *et al*. 2013). To distinguish between these possibilities, we carried out a series of lineage analyses. We first carried out lineage-tracing of glial cells and asked whether they could become neurons. To do this, we used *repo-Gal4* in conjunction with a *flipout-GFP* construct to permanently mark glial lineages. F1 males that were *w[*]; repo-Gal4/P{w[+mC]=Ubi-p63E(FRT.STOP)Stinger}15F2* were injured at within 6 hours of eclosion and aged for 14 days prior to dissection and immunostaining. We observed no *repo à GFP* cells that were also Elav+ (not shown). This indicates that glia do not give rise to neurons through either trans-or dedifferentiation. At first glance, these results appear inconsistent with the observation of EdU+Repo+Elav+ cells shown in **Fig. 2D-D’’’**. However, the cells with hybrid glial-neuronal identity are quite small, and it was reported previously that neuroblasts that undergo reducing divisions are destined for cell death (Siegrist *et al*. 2010). We therefore postulate that the hybrid cells are inviable and do not contribute to the population of new neurons (Class III cells) shown in **Fig. 2**. We note that ∼50% of PH3+ and EdU+ cells post-PTBI are glia (**Fig. 2F**). Thus, although we found no evidence that glia give rise to neurons, glia nonetheless proliferate, especially early post-PTBI.

To test whether neurons can give rise to glia, we used a similar lineage-labeling technique with an *Nsyb-Gal4* driver in a system called Gal4 technique for real-time and clonal expression (G-TRACE) (Evans *et al*. 2009). F1 males that were *w[*];Nsyb-Gal4/P{w[+mC]=UAS-RedStinger}6,P{w[+mC]=UAS-FLP.Exel}3,P{w[+mC]=Ubi-p63E(FRT.STOP)Stinger}15F* were injured within 6 hours of eclosion and aged for 14 days prior to dissection and immunostaining. We observed no *NSyb à GFP* cells that were also Repo+ (not shown). This indicates that neurons do not give rise to new glia post-injury.

To address the possibility that new neurons are created by a quiescent neuroblast-like population, we used a similar lineage labeling technique, this time in combination with a neuroblast driver, *dpn-Gal4*. To ensure that neuroblast cells were not labeled during development, we added a temperature sensitive Gal80 and reared the crosses at 18°C. Under these conditions, the Gal80 prevents transcriptional activation by Gal4, thus keeping the lineage tracing system off. F1 males that were *w[*]; dpn-Gal4/P{w[+mC]=tubP-GAL80[ts]}20; P{w[+mC]=UAS-RedStinger}6,P{w[+mC]=UAS-FLP.Exel}3,P{w[+mC]=Ubi-p63E(FRT.STOP)Stinger}15F* were collected or injured within 6 hours of eclosion and aged for 14 days at 30°C prior to dissection and immunostaining. At 30°C, the temperature sensitive Gal80 protein is inactivated and Gal4 can activate transcription. Indeed, 14 days post-injury, we observed GFP+Elav+ cells in injured brains (arrowheads in **Fig. 4A-A’’’**), but not in uninjured age-matched controls (not shown). These results are consistent with the existence of a quiescent stem cell-like population in the adult *Drosophila* brain that is activated by injury to create new neurons. Several lines of evidence strongly support this view, including the presence of Dpn+ and Ase+ cells near the area of injury 24 hours post-PTBI (**Figs. 3C-D’’’**) and the elevated expression levels *ase, erm,* and *insc* post-PTBI (**Fig. 3A-H**). However, because we do not observe Dpn+ cells in our control uninjured central brains, these putative neuroblast-like cells differ from the neuroblasts present during development in that they lack detectable *dpn* expression until stimulated by PTBI.

**Figure 4.**
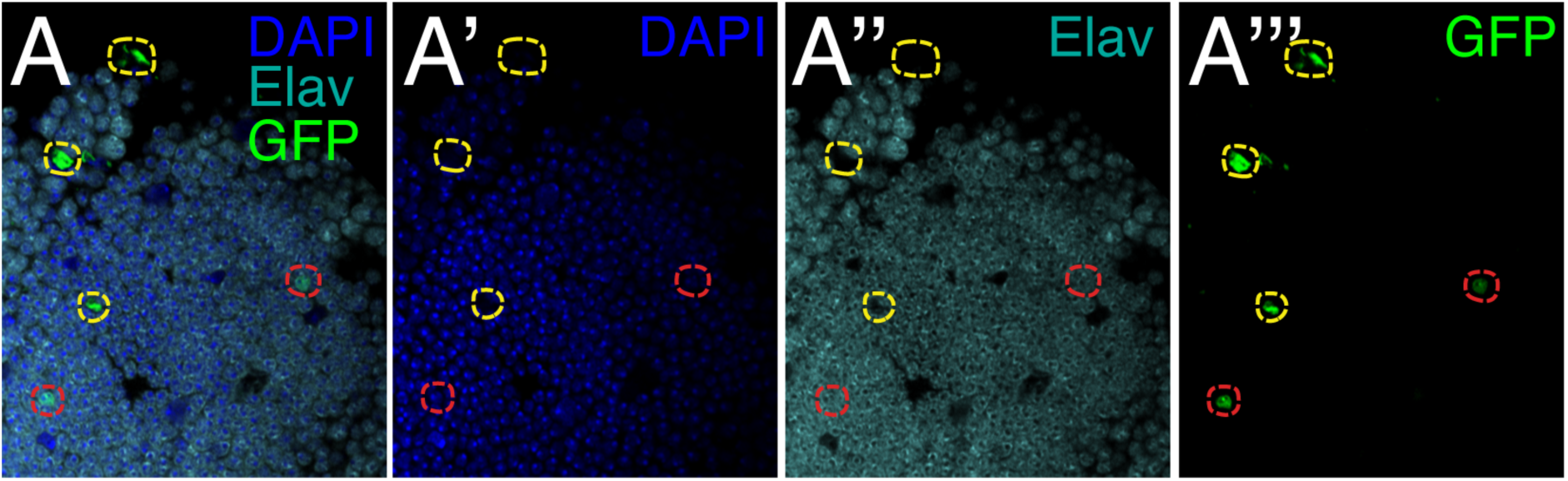
Lineage-tracing demonstrates that new neurons are created by Dpn-expressing cells. Using *dpn-Gal4*, the G-TRACE lineage-tracing system (Evans *et al*. 2009), and a temperature-sensitive Gal80, we lineage-traced *dpn*-expressing cells via their expression of GFP. We observed cells that were GFP+Elav+ (GFP in green; Elav in cyan) at 14 days post-PTBI near the mushroom body (red outlines). The presence of GFP indicates that these cells were either actively expressing or had previously expressed the neuroblast gene *dpn*. GFP+ cells that did not stain with Elav (yellow outlines) also were observed. No GFP+Elav+ cells were observed in age-matched uninjured

### Structural and Functional Regeneration post-PTBI

To visualize new neurons and their projections after PTBI, we utilized perma-twin labeling (Fernandez-Hernandez *et al*. 2013). Perma-twin labeling permanently labels dividing cells and their progeny with either green fluorescent protein (GFP) or red fluorescent protein (RFP). We used adult F1 male flies of the genotype: *w; FRT40A, UAS-CD8-GFP, UAS-CD2-Mir/ FRT40A, UAS-CD2-RFP, UAS-GFP-Mir; act-Gal4 UAS-flp/tub-Gal80^ts^* that were reared at 17°C during development to keep the labeling system switched off. Perma-twin flies were subjected to PTBI within 24 hours of eclosion, and allowed to recover at 30°C for either 2, 7, or 14 days post-injury. At 30°C, the labeling system is active and the progeny of any cells that divide may express either GFP or RFP. As expected, based on our earlier finding that PTBI stimulates cell proliferation, we observed more clones in injured samples than controls at all timepoints (**Fig. 5A-E**). We also found that injured brains had significantly more clones at later timepoints compared to earlier ones, indicating that regeneration and proliferation were progressive and were not limited to the time immediately after the initial injury (**Fig. 5A-E**). Interestingly, we observed large clones at later timepoints that produced new MB neurons (**Fig. A-G**). These new neurons project dendrites correctly to the MB calyx and axons correctly to the MB lobes. This robust regeneration was observed in approximately 50% of the injured brains by 14 days post-PTBI (**Fig. 5N**). Other areas of the brain also grew new neurons and new axon tracts. These include the antennal lobes (AL), the ellipsoid body (EB), and the lateral horn (LH) (**Fig. 5H-M**) in which we observed large clones approximately 26%, 26%, and 20% of the time, respectively (**Fig. 5N**). These data suggest that there is structural repair of the damaged brains.

**Figure 5.**
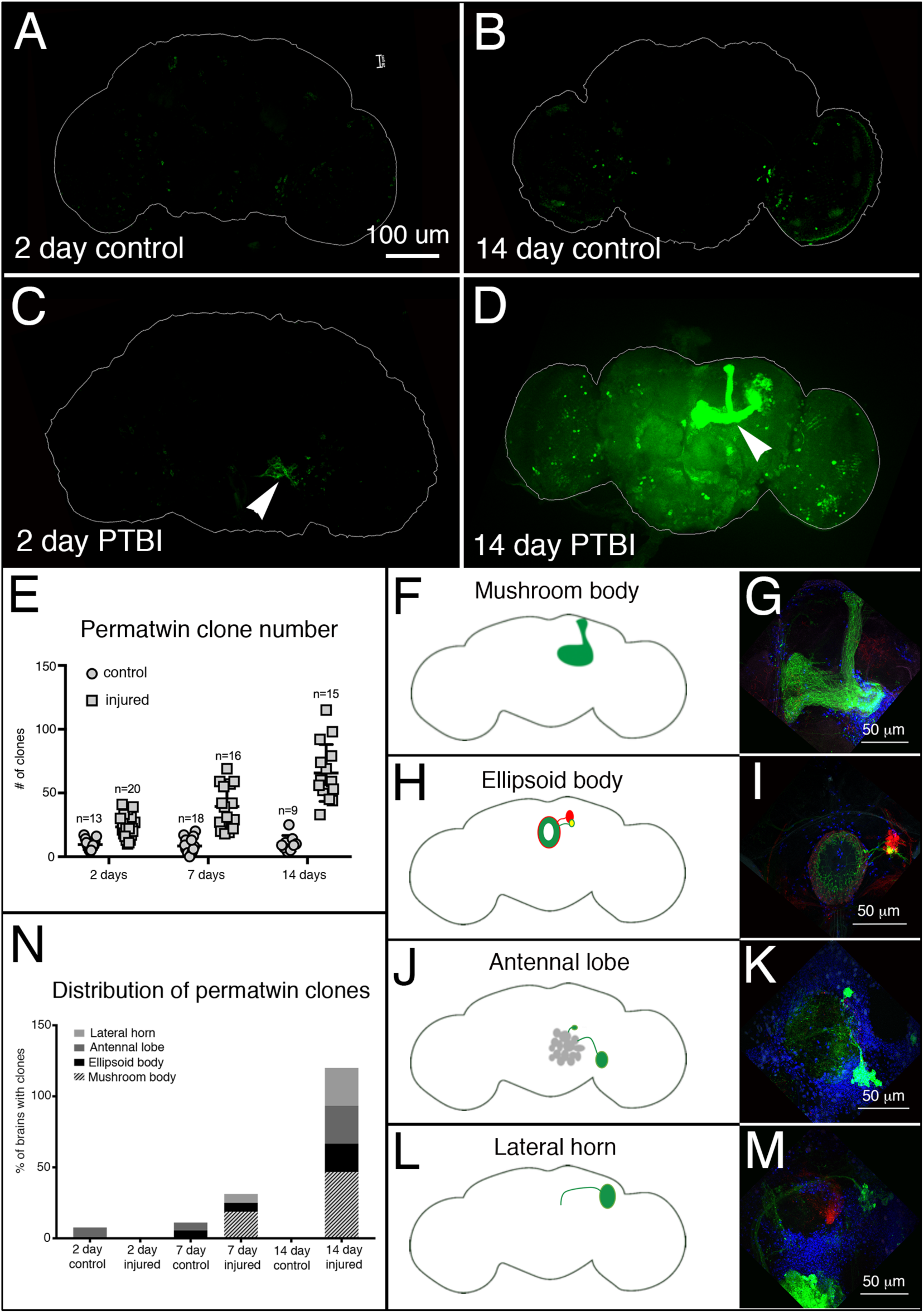
Perma-twin lineage tracing demonstrates brain regeneration and appropriate targeting of axons following PTBI. To analyze neurogenesis after PTBI, we utilized the perma-twin lineage-tracing system (Fernandez-Hernandez *et al*. 2013). This system permanently labels dividing cells and their progeny with either green fluorescent protein (GFP) or red fluorescent protein (RFP). Flies were reared at 17°C to keep the system off during development. Upon eclosion, F1 males carrying perma-twin transgenes were collected, injured and placed at 30°C to recover for either 2 days or 14 days. **A.** In 2-day uninjured controls, there are some GFP+ cells scattered throughout the brain. **B.** At 14 days, there are relatively few GFP+ cells present in the control central brain. **C.** In comparison, 2-day injured brains have more GFP+ cells that tend to cluster near the injury, (arrowhead). **D.** At 14 days post-injury, there are large clones near the site of injury. Some of these clones have axons that project along the mushroom body tracts (arrowhead). Only the GFP channel is shown here; there were similar RFP+ clones in the PTBI samples. **E.** The number of clones increases over time post-PTBI. Control uninjured brains (n=13) have an average of 10 clones at 2 days while 2-day PTBI brains (n=20) have an average of 23 clones (p<0.00002). At 7 days, control brains had an average of 9 clones per brain (n=18), while 7-day PTBI brains had an average of 39 clones per brain (n=16) (p-value<0.00000002). This is significantly more than the number of clones seen at 2 days post-injury (p-value<0.0009). In 14-day control brains, there are an average of 10 clones per brain, which is not significantly different from the 2-day and 7-day controls. However, at 14 days post-PTBI, there are an average of 66 GFP+ clones, which is significantly more than either age-matched controls (p<0.0000003) or 2-day post-PTBI brains (p-value<0.0001). Error bars reflect SD. **F-M.** PTBI stimulates clone formation in multiple regions in the brain. Panels on the left side are schematics of brain regions where large clones were found 14 days post-PTBI (**A**, **H**, **J**, **L**). Panels on the right show high magnifications of representative brains (**G**, **I**, **K**, **M**). Many 14-day brains had clones that projected to particular target areas. These included the mushroom body (MB) (**F**, **G**), the ellipsoid body (EB) (**H**, **I**), the antennal lobe (AL) (**J**, **K**), and the lateral horn (LH) (**L**, **M**). **N.** Both clone number and clone size increase with time post-PTBI. The proportions of brain regions with large clones were calculated at 2, 7, and 14 days in controls and injured brains. At 2 days, approximately 8% of control brains (n=13) showed AL clones, while in 2-day injured brains (n=20), there were no AL clones. In 7-day control brains (n=18), 6% had AL and 6% had EB clones. At 7 days post-PTBI (n=16), 6% of brains also had AL clones, 6% had EB clones, and 19% had large MB clones. At 14 days, control brains (n=9) did not exhibit any specific areas with clones, while 47% of PTBI brains (n=15) had MB clones, 20% of PTBI brains had AL clones, and 27% of PTBI brains had EB clones, and 27% had LH clones.

In order to assess functional recovery post-PTBI, we asked whether *Drosophila* locomotor function is impaired by PTBI and, if so, whether function is restored at later timepoints. 2-day post-PTBI flies, exhibited significantly different locomotor profiles from the stereotypic locomotory patterns of age-matched controls (**Fig. 6A**). However, by 14 days post-injury, by which time we also observed the generation of new neurons and axon tracts, the injured flies displayed comparable locomotor profiles to age-matched controls (**Fig. 6B**). These data indicate that PTBI significantly impacts motor function, and that this damage is largely repaired by 14 days.

**Figure 6.**
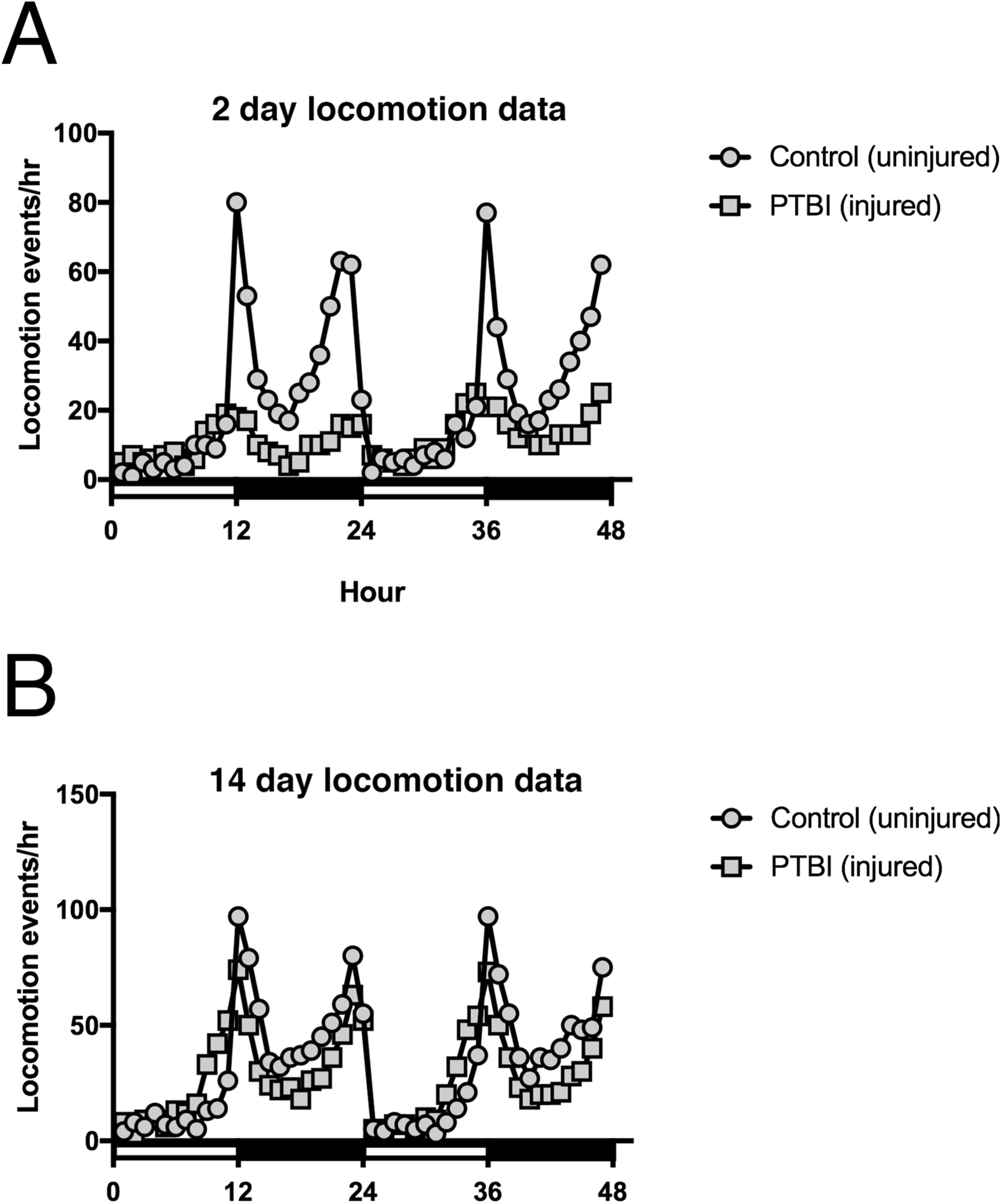
Locomotor defects observed at 2 days post-PTBI are reversed by 14 days post-PTBI. To assay for functional recovery post-PTBI, we examined locomotor function. The 2 and 14 day and age-matched uninjured controls were placed in the *Drosophila* Activity Monitor (DAM) system (TriKinetics, Waltham, MA) to record locomotory behavior. For each condition, 32 newly eclosed males were collected, either injured or kept as uninjured controls, and individually monitored over a 48 hour window. **A.** 2-day control uninjured flies displayed stereotypic locomotory patterns throughout a 24-hour period. However, 2-day post-PTBI flies, exhibited significantly different locomotor profiles (p-value<0.001). **B.** Nonetheless, by 14 days, PTBI flies display comparable locomotor profiles to age-matched controls.

## Discussion

Following a penetrating traumatic brain injury (PTBI), we find that the adult *Drosophila* central brain has regenerative potential (**Fig. 7**). We demonstrate that PTBI rapidly stimulates cell proliferation (**Fig. 1**) and that a robust proliferative response occurs primarily in young adult flies and diminishes with age (**Figs. S1A and S1B**). Our data also indicate that age plays an important role in the adult *Drosophila*’s ability to survive a traumatic injury (**Fig. S2B**), consistent with the age-dependence of PTBI survival reported by Sanuki *et al*. (Sanuki *et al*. 2019). Within one week of PTBI, both new glia and new neurons have been created (**Fig. 2**). At early timepoints post-PTBI, but not in control brains, we observe dividing neuroblast-like cells that are Dpn+ (**Fig. 3**). Other neural progenitor genes such as *ase*, *insc,* and *erm* exhibit elevated transcript levels at early timepoints post-injury (**Fig. 3**). Using cell lineage-tracing techniques, we found that new neurons are generated by cells that had expressed *dpn* (**Fig. 4**). These *dpn*-expressing cells were not observed in uninjured controls. The newly created neurons contribute to the overall regeneration of damaged brain tissue, particularly near the mushroom body (**Fig. 5**). The timing of regeneration corresponds with the timing of recovery of the locomotor activity disrupted by PTBI (**Fig. 6**). This supports the idea that the newly generated neurons are both functional and properly connected. However, we cannot rule out the possibility that other parts of the brain are compensating for the damaged areas and that the behavioral improvement is not a direct consequence of the new neurons.

**Figure 7.**
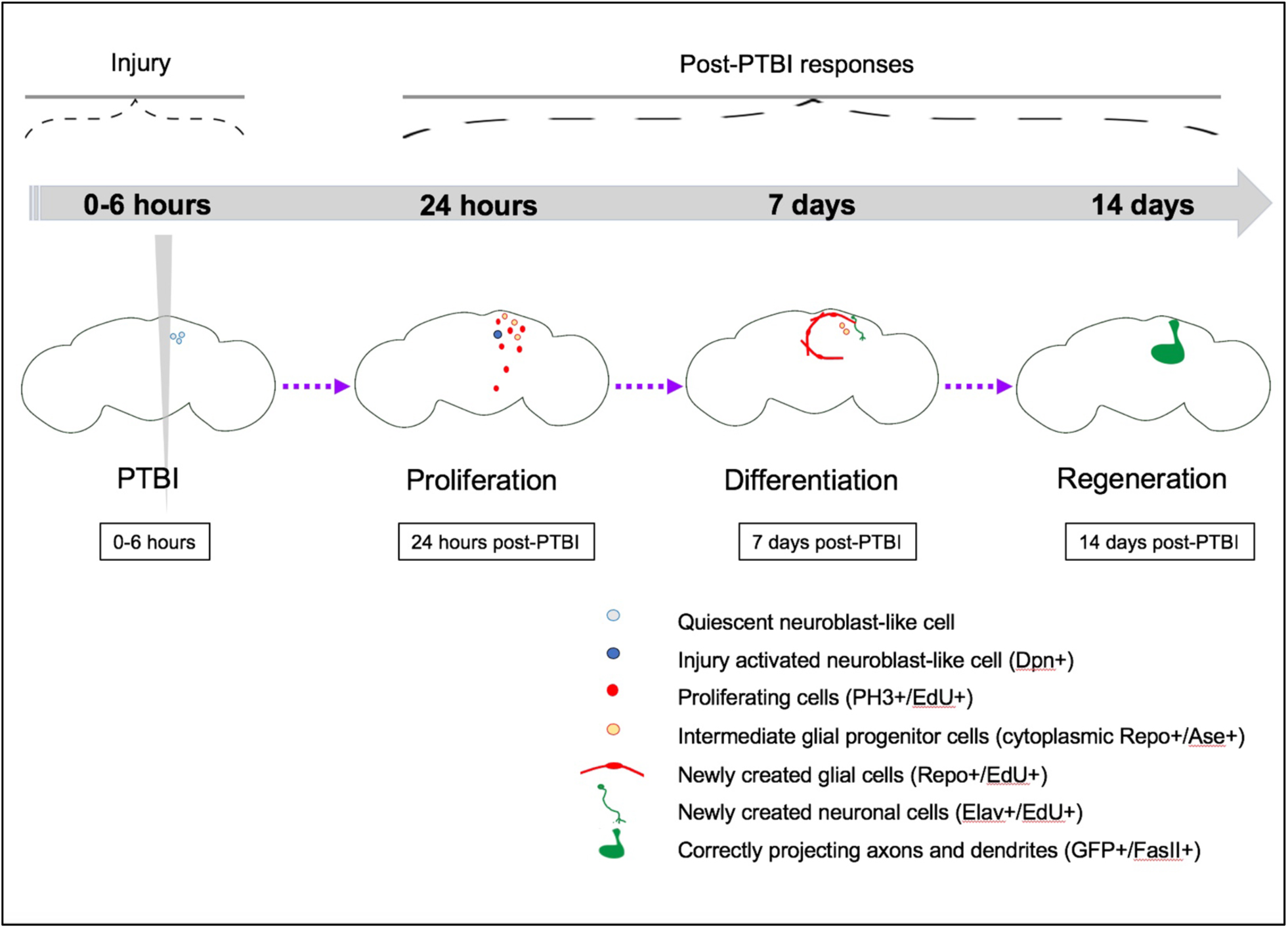
Summary model for regeneration following penetrating traumatic brain injury (PTBI). We propose that in young adult *Drosophila* there are quiescent NB-like cells within the central brain that lack expression of canonical NB genes. By 24 hours post-PTBI, the quiescent NB-like cells are activated, express NB genes, and have begun to proliferate. At both 4 hours and 24 hours post-PTBI, there is a wave of cell death as assayed using TUNEL. At 7 days, the proliferation rate is still high, and many of the new cells have adopted mature cell identities, becoming neurons or glia. At 10 days post-PTBI, there is no longer a difference in TUNEL+ cells between uninjured brains and injured brains, indicating that the wave of cell death has ended. Because the peaks of both cell death and proliferation occur at the same time post-injury, this could explain why there is not a significant increase in the number of EdU+ cells seen at 7 days compared to the number of PH3+ cells seen at 24 hours. However, by 10 days, cell death is back to control levels while proliferation has decreased but is still slightly above baseline levels. This could explain why there is an increase in the number of EdU+ between 7 and 14 days. At 14 days post-PTBI, there are large clones of new neurons with axons and dendrites correctly projecting to their respective target areas. Locomotor defects are also restored by 14 days, suggesting that adult *Drosophila* are able to regenerate functionally as well as structurally.

We were intrigued to observe cells of hybrid glial (Repo+) and neuronal (Elav+) identity in injured brains (**Fig. 2**), and initially thought these might represent a transitional state from glia to neuron or from glia to neural precursor. Repo+Elav+ cells are not present in uninjured control adult brains, but have been reported during larval development (Berger *et al*. 2007) and in certain brain tumors (Beaucher *et al*. 2007). Elav also is known to be transiently expressed in some neuroblast-like cells during development (Beaucher *et al*. 2007). Thus, the presence of dividing cells that are Repo+Elav+ is consistent with a less differentiated state. However, our lineage studies do not support the hypothesis that these hybrid cells are the progenitors of either glia or neurons. Specifically, while glia give rise to new glia, lineage tracing did not reveal any new neurons derived from cells with prior glial identity, nor did it reveal any new glia derived from cells with prior neuronal identity. Also notable is that the cells possessing the hybrid glial and neuronal fate are quite small. Reducing divisions and small cell size previously were correlated with apoptosis in Drosophila brains (Siegrist *et al*. 2010). Together, these results are most consistent with the idea that cells possessing the hybrid glial and neuronal fate are inviable.

Previous studies have indicated a higher occurrence of certain brain tumors, such as glioblastomas, in people that have previously experienced TBI (Tyagi *et al*. 2016). However, following PTBI, we do not find evidence of unregulated cell division. When young adult flies are subjected to PTBI, there is a robust proliferative response that diminishes with time post-PTBI (**Fig. S1A**). The proliferative response also diminishes with age and is negligible by 2 weeks post-eclosion (**Fig. S1B**). By histology, we also do not see any evidence of tumor formation by 25 days post-PTBI (**Fig. S2E,G**). Neurodegeneration is another common consequence of brain injury both in mammals (reviewed in (Graham and Sharp 2019)) and Drosophila (Katzenberger *et al*. 2013) and is thought to result from secondary brain injury, i.e. from cascades triggered by trauma and not the trauma itself (reviewed in (Ng and Lee 2019)). We do observe limited neurodegeneration following PTBI (**Fig. S2E,G**). This is consistent with a small amount of cell death in the first hours after PTBI (**Fig. S2C-D’, F**). However, cell death is restricted by 24 hours post-PTBI and indistinguishable from controls by 10 days post-PTBI (**Fig. S2F**). This limited cell death and neurodegeneration may contribute to a small reduction in lifespan observed post-PTBI (**Fig. S1A**). Together, our results support the ideas that secondary injury is minimal in our PTBI model and that PTBI does not stimulate uncontrolled cell division. The mechanisms underlying what appears to be a tightly-regulated process need to be further analyzed to understand how *Drosophila* are able to give a measured proliferative cell response to regenerate damaged tissue.

We detect both new glia and new neurons post-PTBI, but these cell types are not generated in equal proportions or on the same time scale. There is an initial wave of gliogenesis, followed by a delayed wave of neurogenesis (**Fig. 2**). Glial proliferation is a known response to neuronal injury in both mammals and Drosophila, with the added glia participating in phagocytosis of cellular debris (reviewed in (Burda and Sofroniew 2014; Pekny and Pekna 2016)). The newly-generated glia may play similar roles following PTBI.

In addition to proliferating glial cells, we have identified a distinct population of cells that divide to give rise to new neurons. These cells do not express either Repo nor Elav, i.e. they are neither glia nor neurons. In addition, although our lineage analysis indicates that the new neurons arise from cells that express *dpn* (**Fig. 4**), we do not detect *dpn* expression in uninjured adult central brains with either Dpn antibodies or *dpn* reporters. We therefore propose that *dpn* expression is activated by central brain injury, and that cells that activate *dpn* expression give rise to new neurons. *dpn* encodes a basic helix-loop-helix (bHLH) transcriptional repressor orthologous to mammalian HES1 (Bier *et al*. 1992; Younger-Shepherd *et al*. 1992). During normal Drosophila neural development Dpn represses the expression of neural differentiation genes (Kang and Reichert 2015; Ramon-Canellas *et al*. 2019; Li and Hidalgo 2020). Thus, the upregulation of *dpn* expression post-PTBI may confer neural progenitor fate on these cells and be an essential step in adult neurogenesis. Our data support the idea that there is a quiescent population of cells in uninjured brains that cannot be detected with standard neuronal, glial, or neuroblast markers, but that nonetheless has regenerative potential. If such cells exist in *Drosophila*, perhaps they also exist in humans. An important avenue for future research includes the generation of gene expression profiles and markers for these progenitor cells such that they can be visualized in and sorted from uninjured brains in order to explore the mechanisms of their activation.

We note that in the optic lobe, cells with cytoplasmic Dpn protein were identified and proposed to be quiescent neural progenitors, with nuclear translocation of Dpn following injury correlated with adult neurogenesis (Fernandez-Hernandez *et al*. 2013). We do not observe either nuclear or cytoplasmic Dpn protein in the uninjured central brain. Nor have we observed GFP expression driven by the *dpn-GAL4* reporter in the uninjured central brain. Thus, while there is a quiescent neural progenitor cells identifiable with standard markers in the optic lobe, this does not appear to be the case in the central brain. A second difference between adult neurogenesis in the optic lobe and the central brain involves expression of the neural progenitor gene *ase*. In the optic lobe, *ase* is not upregulated by injury (Fernandez-Hernandez *et al*. 2013). In contrast, in the central brain after PTBI, there is *ase* upregulation as assayed by both qRT-PCR and immunohistochemistry (**Fig. 3**). Interestingly, the *ase*-expressing cells are in close proximity to *dpn*-expressing cells (**Fig. 3**). This is reminiscent of normal neural development in Type II neuroblast lineages (Bello *et al*. 2008; Boone and Doe 2008; Bowman *et al*. 2008). Nonetheless, the expression of both Type I (*insc*) and Type II (*erm*) lineage attributes suggests that these cells do not recapitulate larval neurogenesis. A third difference between adult neurogenesis in the optic lobe and the central brain is that optic lobe injury does not result in the generation of new glia. In contrast, glial proliferation is rapid and robust following central brain injury (**Fig. 2**). Together, these results indicate that adult central brain neurogenesis differs from adult neurogenesis in the optic lobe. Given that the neural progenitors of the optic lobes and central brain follow distinct developmental trajectories (Ramon-Canellas *et al*. 2019), it perhaps should not come as a surprise that their regenerative capacities and programs also are different.

Using a PTBI paradigm, we have been able to establish that young adult *Drosophila* are capable of robust regeneration, with the creation of new neurons and glia and functional recovery from locomotor defects by 14 days post-PTBI. Further questions remain about the origin and properties of the neural progenitor cells and the molecular mechanisms that trigger regeneration. Nonetheless, because brain regeneration can be stimulated by mild injury in adult Drosophila, there is an avenue by which to identify these cells and mechanisms for future study. In addition, the Drosophila adult central brain now provides a novel model system for screening pharmacologic agents for those that activate the regenerative program. This could lead to new therapeutic approaches for both neurodegenerative diseases and brain injuries.

## Materials & Methods

### Fly Stocks and Rearing

Unless otherwise specified, flies were reared at 25°C on a standard cornmeal-sugar medium. The following stocks were obtained from the Bloomington Drosophila Stock Center (BDSC): **#854** (*w[*]; P{w[+mW.hs]=GawB}OK107 ey[OK107]/ln(4)ci[D], ci[D] pan[ciD] sv[spa-pol]*; **#1495** (*y[1] w[1]*); **#4539** *y[1] w[*]; P{w[+mC]=UAS-FLP.D}JD1*; **#5130** (*y[1] w[1]; Pin[Yt]/CyO; P{w[+mC]=UAS-mCD8::GFP.L}LL6)*; **#7018** (*w[*]; sna[Sco]/CyO;P{w[+mC]=tubP-GAL80[ts]}ncd[GAL80ts-7]*); **#7019** (*w[*];P{w[+mC]=tubP-GAL80[ts]}20;TM2/TM6B, Tb[1]*; **#7415** (*w^1118^; P{w[+m*]=GAL4}repo/TM3, Sb^1^*); **#28281** (*w[*];P{w[+mC]=UAS-RedStinger}6,P{w[+mC]=UAS-FLP.Exel}3,P{w[+mC]=Ubi-p63E(FRT.STOP)Stinger}15F*); **#32251** (*w[*]; P{w[+mC]=Ubi-p63E(FRT.STOP)Stinger}15F2*); **#47859** *w[1118]; P{y[+t7.7] w[+mC]=GMR13CO2-GAL4}attP2*; **#51635** (*y^1^ w*; P{w[+m*]=nSyb-GAL4S}3*): **#65408** (*P{w[+mC]=UAS-Stinger}2, P{w[+mC]=UAS-hid.Z}2/CyO*). Other lines used were *ase-Gal4/CyO; Dr/TM6B* (a gift of Dr. Cheng-Yu Lee); *w; FRT40A, UAS-CD8-GFP, UAS-CD2-Mir; act-Gal4 UAS-flp/TM6B*; and *w; FRT40A, UAS-CD2-RFP, UAS-GFP-Mir; tub-Gal80ts/TM6B* (both gifts of Dr. Eduardo Moreno).

### Standard cross

To minimize potentially confounding genetic background effects and differences in results due to sex and temperature, we routinely analyzed F1 males of a standard genotype. This standard genotype results from an outcross between a homozygous strain expressing green fluorescent protein (GFP) in the mushroom body (*w*; UAS-mCD8-GFP;; OK107-Gal4*) and the widely used laboratory strain *y^1^ w^1^*. This cross was maintained at 25°C. PTBI flies were kept at 25°C until their brains were dissected and fixed for analysis. Unless otherwise specified, brains of young adult male *Drosophila* generated by this outcross were injured within 6 hours of eclosion.

### Perma-twin flies

Perma-twin flies were generated by crossing *w; FRT40A, UAS-CD2-RFP, UAS-GFP-Mir; tub-Gal80^ts^/TM6B* virgin females to *w; FRT40A, UAS-CD8-GFP, UAS-CD2-Mir; act-Gal4 UAS-flp/TM6B* males (Fernandez-Hernandez *et al*. 2013). These crosses were maintained at 17°C. F1 progeny of the genotype: *w; FRT40A, UAS-CD8-GFP, UAS-CD2-Mir/ FRT40A, UAS-CD2-RFP, UAS-GFP-Mir; act-Gal4 UAS-flp/tub-Gal80^ts^* were collected at eclosion, subjected to PTBI or kept as uninjured controls and maintained at 30°C until their brains were dissected and fixed for analysis.

### G-TRACE crosses

Lineage-labeling was accomplished using a G-TRACE line (**#28281** (*w[*];P{w[+mC]=UAS-RedStinger}6,P{w[+mC]=UAS-FLP.Exel}3,P{w[+mC]=Ubi-p63E(FRT.STOP)Stinger}15F*) (Evans *et al*. 2009) crossed to various Gal4 driver strains listed above. These crosses were maintained at 17°C. F1 progeny of the desired genotyped were selected at eclosion, subjected to PTBI or kept as uninjured controls and maintained at 30°C for 14 days when their brains were dissected and fixed for analysis.

### Penetrating Traumatic Brain Injury

To induce PTBI, we used thin metal needles (∼12.5 μm diameter tip, 100 μm diameter rod; Fine Science Tools) sterilized in 70% ethanol to penetrate the head capsule of CO_2_-anesthetized adult flies. Injured flies were transferred back to our standard sugar food for recovery and aging. For immunohistochemical analyses, we unilaterally injured brains on their right sides. For qRT-PCR analysis, to amplify the molecular responses, brains were injured bilaterally.

### Immunohistochemistry

Brains were dissected in PBS (phosphate-buffered saline; 100 mM K_2_HPO4, 140 mM NaCl pH 7.0) and fixed in a 3.7% formaldehyde in a PEM (100 mM PIPES, 2 mM EGTA, 1 mM MgSO_4_) solution for 20 minutes at 25°C. Fixed brain samples were washed in PT (PBS and 0.1% Triton X-100), blocked with 2% BSA in PT solution (PBT), and then incubated with primary antibodies overnight at 4°C in PBT. Following primary antibody incubation, the samples were washed with PT (5 times over the course of an hour) and incubated overnight in secondary antibody at 4°C. The next day, samples were washed in PT, stained with DAPI (1:10,000, ThermoFisher) for 8 minutes, and mounted in Vectashield anti-fade mountant (Vector Labs) and imaged using a Nikon A1RS system and analyzed using the Nikon NIS Elements software. Cell counting was done both manually and using the Nikon NIS-Elements software to analyze regions of interest (ROIs) with a threshold of over 1000 and an area of at least 10μm.

The primary antibodies used in this study were: rabbit anti-PH3 (1:500, Santa Cruz Biotechnology, Inc); mouse anti-FasII (1:20, Developmental Studies Hybridoma Bank; DSHB); mouse anti-Repo (1:20, DSHB); rat anti-Elav (1:20, DSHB); mouse anti-Pros (1:20, DSHB); and rat anti-Dpn (1:50, AbCam). Secondary antibodies used were: anti-rabbit Alexa Fluor 568 (1:200, ThermoFisher); anti-rabbit Cy5 (1:400, Jackson ImmunoResearch, Inc.); anti-mouse Cy5 (1:100, Jackson ImmunoResearch, Inc.); anti-rat Alexa Fluor 488 (1:400, ThermoFisher); anti-rat Alexa Fluor 568 (1:400, ThermoFisher); and anti-rat Cy5 (1:200, Jackson ImmunoResearch, Inc.).

### EdU labeling

The standard injury method was used on flies for 5-ethynyl-2’-deoxyuridine (EdU) labeling, except flies were fed 50 mM EdU in 10% sucrose solution on a size 3 Whatman filter for six hours prior to PTBI and allowed to recover on the same solution for the desired amount of time. The EdU solution was replaced every 24 hours. Brains were dissected, processed, and antibody stained as described above with the exception of using buffers without azide. To detect EdU incorporation, Click-IT® reagents from InVitrogen were used according to the manufacturer’s instructions. The brains then were antibody stained mounted and imaged as described above.

### Quantitative Real-Time PCR

Transcript levels of target genes were measured by quantitative real-time PCR (qRT-PCR) using methods described in (Ihry *et al*. 2012). RNA was isolated from appropriately staged animals using TRIzol Reagent used according to the manufacturer’s instructions (Thermo Fisher Scientific). cDNA was synthesized from 40 to 400 ng of total RNA using the SuperScript III First-Strand Synthesis System (Invitrogen). qPCR was performed on a Roche 480 LightCycler using the LightCycler 480 DNA SYBR Green I Master kit (Roche). In all cases, samples were run simultaneously with three independent biological replicates for each target gene, and *rp49* was used as the reference gene. To calculate changes in relative expression, the Relative Expression Software Tool was used (Pfaffl *et al*. 2002). We used the following primers to detect transcript levels: *ase* Forward: 5’-CAGTGATCTCCTGCCTAGTTTG-3’ & Reverse: 5’-GTGTTGGTTCCTGGTATTCTGATG-3’ (gift from Stanislava Chtarbanova); *dpn* Forward: 5’-CGCTATGTAAGCCAAATGGATGG-3’ & Reverse: 5’-CTATTGGCACACTGGTTAAGATGG-3’ (gift from Stanislava Chtarbanova); *elav* Forward: 5’-CGCAGCCCAATACGAATGG-3’ & Reverse: 5’-CATTGTTTGCGGCAA GTAGTTG-3’ (Fly Primer Bank; (Hu *et al*. 2013)); *erm* Forward: 5’-GTCCCCTAAAGTTTTCGATAGCC-3’ & Reverse: 5’-GAGTCATAGTTGACAGTGGATGG-3’ (Fly Primer Bank); *insc* Forward: 5’-CCCTGGGCAATCTGTCCTG-3’ & Reverse: 5’-GAGAAGCCCGAATCCTGACT-3’ (Fly Primer Bank); *myc* Forward: 5’-AGCCAGAGATCCGCAACATC-3’ & Reverse: 5’-CGCGCTGTAGAGATTCGTAGAG-3’ (Fly Primer Bank); *repo* Forward: 5’-TCGCCCAACTATGTGACCAAG-3’ & Reverse: 5’-CGGCGCACTAATGTACTCG-3’ (Fly Primer Bank;; *Rp49* Forward: 5’-CCAGTCGGATCGATATGCTAA-3’ & Reverse: 5’-ACGTTGTGCACCAGGAACTT-3’ (Ihry *et al*. 2012).

### Locomotor assays

0-6-hour post-eclosion *OK107/yw* males were collected, subjected to PTBI, and aged to 2 days and 14 days, respectively. The 2 and 14 day injured and age-matched uninjured controls were placed in the *Drosophila* Activity Monitor (DAM) system (TriKinetics, Waltham, MA) to record locomotory behavior. The circadian locomotor activity of flies was assayed and analyzed as previously described (Hamblen *et al*. 1986; Sehgal *et al*. 1992). For each experiment and condition, 32 flies were individually analyzed.

### Statistical analysis

For all cell/clone counting and locomotor assays, counts were expressed as means ± standard deviations. Two-tailed t-tests were performed using GraphPad Prism Version 8.3.0 for Mac (GraphPad Software, La Jolla California USA, www.graphpad.com). An alpha value of 0.05 was considered significant. The following symbols represent significance; * significant at p ≤ 0.05; ** significant at p ≤ 0.01; *** significant at p ≤ 0.001; **** significant at p ≤ 0.0001.

### Reagent sharing statement

Strains are available upon request. The authors affirm that all data necessary for confirming the conclusions of the article are present within the article, figures, and tables.

## Acknowledgements

We are grateful to Becky Katzenberger and Sarah Neuman for technical assistance; Eduardo Moreno for sharing the Perma-twin stocks; and Stanislava Chtarbanova for the *ase* and *dpn* primers. We also would like to thank Barry Ganetzky and David Wassarman for lively discussions that undoubtedly improved the science and Kent Mok, Cayla Guerra, Bailey Spiegelberg, and Shawn Ahern-Djamali for their contributions to the laboratory. The FasII, Elav and Repo monoclonal antibodies were obtained from the Developmental Studies Hybridoma Bank, created by the NICHD of the NIH and maintained at The University of Iowa, Department of Biology, Iowa City, IA 52242. Most of the Drosophila strains used in this study were obtained from the Bloomington Drosophila Stock Center (BDSC; NIH P40OD018537). This work was supported by NIH T32 GM007133 (KLC and KM); NIH NS090190 (GBF); NIH NS102698 (GBF); the University of Wisconsin Graduate School (GBF); and the Women in Science and Engineering Leadership Institute (WISELI) (GBF).

## Supplemental Figure Legends

**Supplemental Figure 1.**
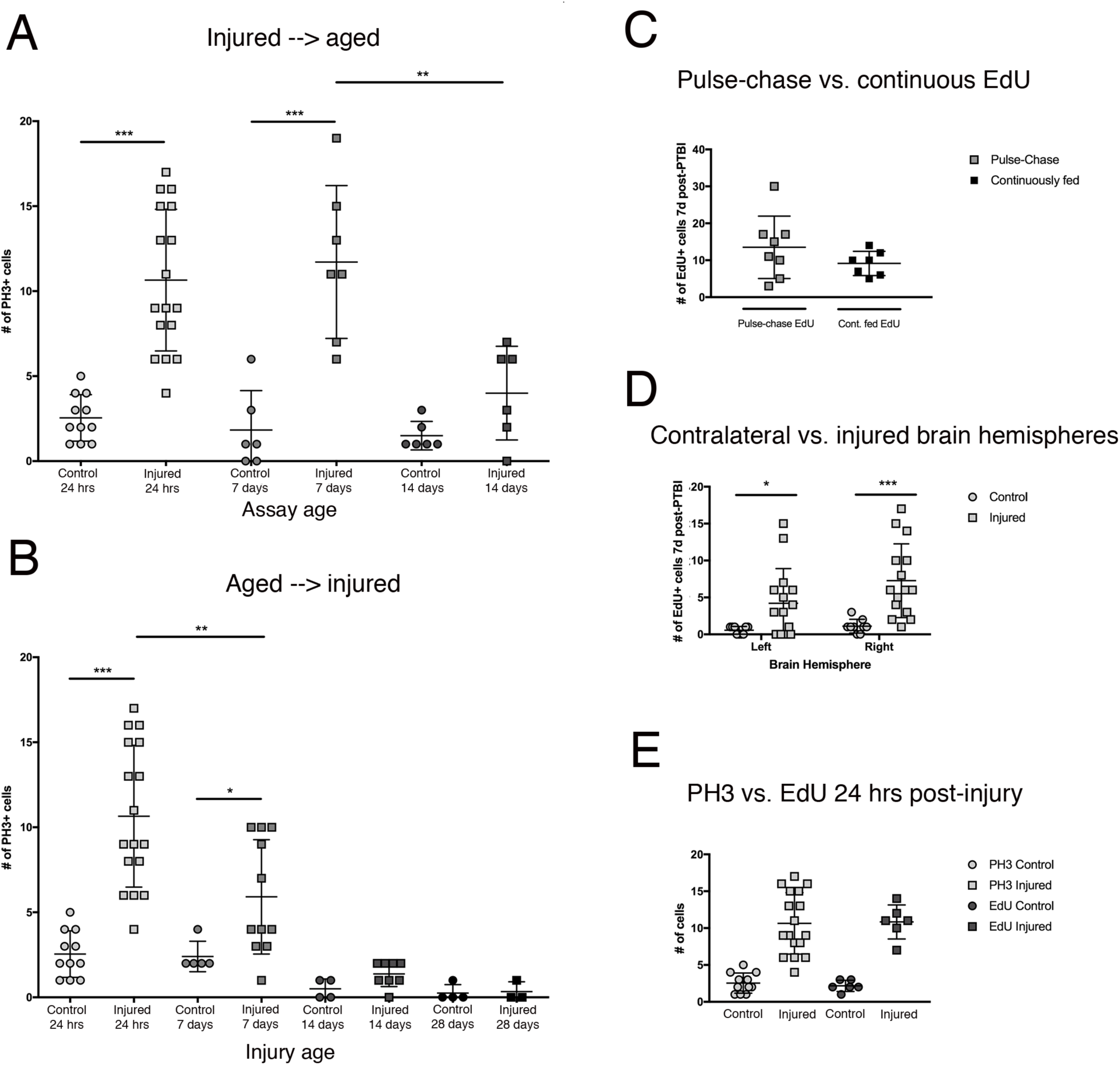
**S1A. The number of actively dividing cells decreases with time post-injury.** In Fig. 1, we utlilized EdU incorporation at 7 and 14 days to measure cell proliferation. However, EdU labeling is cumulative, allowing one to identify all cells that divided during the labeling period. To obtain snapshots of proliferation in real time, we also carried out PH3 labeling at these same time points post-PTBI. For this experiment, we collected newly eclosed male flies, subjected them to PTBI, and aged them for the specified time periods. We find similar numbers of proliferating cells at 24 hours and 7 days post-PTBI, and observe a strong reduction in proliferation at 14 days post-PTBI. At 24 hours, control brains have an average of 3 PH3+ cells (n=11 brains, 28 cells), while PTBI brains have an average of 11 PH3+ cells (n=17 brains, 181 cells). At 7 days, control brains have an average of 2 PH3+ cells (n=6 brains, 11 cells), while PTBI brains have an average of 12 PH3+ cells (n=7 brains, 82 cells). At 14 days, control brains have an average of 2 PH3+ cells (n=6 brains, 9 cells), while PTBI brains have an average of 4 PH3+ cells (n=6 brains, 24 cells). Unpaired t tests for PTBI to control comparisons at these 3 times points are p<0.0001, p<0.0005, and p<0.06, respectively. **S1B. Robust proliferative responses decrease with age.** To explore whether age impacts the amount of cell proliferation that occurs post-injury, we compared newly eclosed adult males to animals aged to 7 days, 14 days, and 28 days prior to PTBI. We then used anti-PH3 to assay cell proliferation 24 hours after injury. Flies injured at within 6 hours of eclosion had an average of 11 PH3+ cells/brain (n=17 brains, 182 cells) compared to an average of 3 PH3+ cells/brain in age-matched controls (n=11 brains, 28 cells). Flies that were aged to 7 days, then subjected to PTBI had an average of 6 PH3+ cells/brain (n=11 brains, 65 cells) compared to age-matched controls which had an average of 2 PH3+ cells/brain (n= 5 brains, 12 cells). When flies were aged to 14 days prior to PTBI, and assayed 24 hours later, there was an average of 1 PH3+ cell/brain (n=8 brains, 11 cells) similar to age-matched controls which also averaged 1 PH3+ cell/brain (n=4 brains, 2 cells). 28-day uninjured control (n=4, 1 cell) and PTBI (n=3, 1 cell) flies both averaged 0 PH3+cells/brain. Unpaired t tests for PTBI to control comparisons at these 4 time points are p<0.0001, p<0.04, p<0.07, and p<0.84, respectively. **S1C. Pulse-chase and continuously fed EdU animals have similar numbers of EdU+ cells at 7 days post-PTBI.** To assess whether mitotically active cells survive post-PTBI, we used two methods of feeding EdU, pulse-chase (flies are fed EdU for 4 days, then placed on standard sugar food for 3 days) and continuously fed (flies are fed EdU every day before being assayed). At 7 days post-PTBI, we find that pulse-chase PTBI brains have an average of 14 EdU+ cells (n=8, 108 cells) while continuously fed PTBI brains have an average of 9 EdU+ cells (n=7, 64 cells). Although there is a trend of fewer EdU+ cells in the continuously fed animals, an unpaired t test reveals that this is not significantly different (p<0.22). Error bars reflect the standard deviation (SD). **S1D. Cell proliferation is not limited to the damaged right hemisphere.** In order to determine where cell proliferation occurs post-PTBI, we counted EdU+ cells in the left and right hemispheres of the central brain. At 7 days, the left hemispheres of control brains have an average of 1 EdU+ cells (n=9 brains, 5 cells) while PTBI brains have an average of 4 EdU+ cells (n=15 brains, 63 cells; p<0.04). In the right hemisphere, control brains have an average of 1 EdU+ cell (n=9 brains, 10 cells), while PTBI brains have an average of 7 EdU+ cells (n=15 brains, 109 cells; p<0.002). Error bars reflect the standard deviation (SD). **S1E. There is no significant difference in the number of PH3+ and EdU+ cells observed 24 hours post-PTBI.** We use two methods, anti-PH3 immunochemistry to label mitotic cells and EdU labeling of newly synthesized DNA to assay cell division. To assess the extent to which anti-PH3 and EdU labeling are comparable, we evaluated both control and PTBI brains with both methods at 24 hours. We expected to detect more EdU-labeled cells than anti-PH3 labeled cells post-PTBI because anti-PH3 transiently labels cells during M phase of the cell cycle while EdU labeling is cumulative. Instead, we observed similar numbers of labeled cells with the two assays. Specifically, in control brains, there were an average of 3 PH3+ cells (n=11 brains, 28 cells) and 2 EdU+ cells (n=6 brains, 13 cells). In PTBI brains, there were an average of 11 PH3+ cells (n=17 brains, 181 cells) and 11 EdU+ cells (n=6 brains, 65 cells). Thus, while control and injured brains displayed significant differences in cell proliferation with both assays (PH3: p<0.0001, EdU: p<0.0005), the number of proliferating cells detected with the two methods was not significantly different. Error bars reflect the standard deviation (SD). There are several potential explanations for this result. For instance, it could be that EdU does not efficiently penetrate the blood brain barrier and/or diffuse through the brain tissue. In this case, only a subset of proliferating cells would be labeled, and EdU labeling would underreport cell proliferation. Alternatively, some of the dividing cells may die. Indeed, DNA synthesis does precede cell death in some contexts (e.g. (Rimkus *et al*. 2008)). However, the fact that there are similar numbers of EdU positive cells in post-PTBI brains when animals are continuously fed EdU and when they are pulse-chased with EdU (**Fig. S1C**), supports the idea that most dividing cells are viable. It therefore seems likely that EdU does not efficiently penetrate the blood brain barrier and/or diffuse through the brain tissue, particularly in older animals.

**Supplemental Figure 2.**
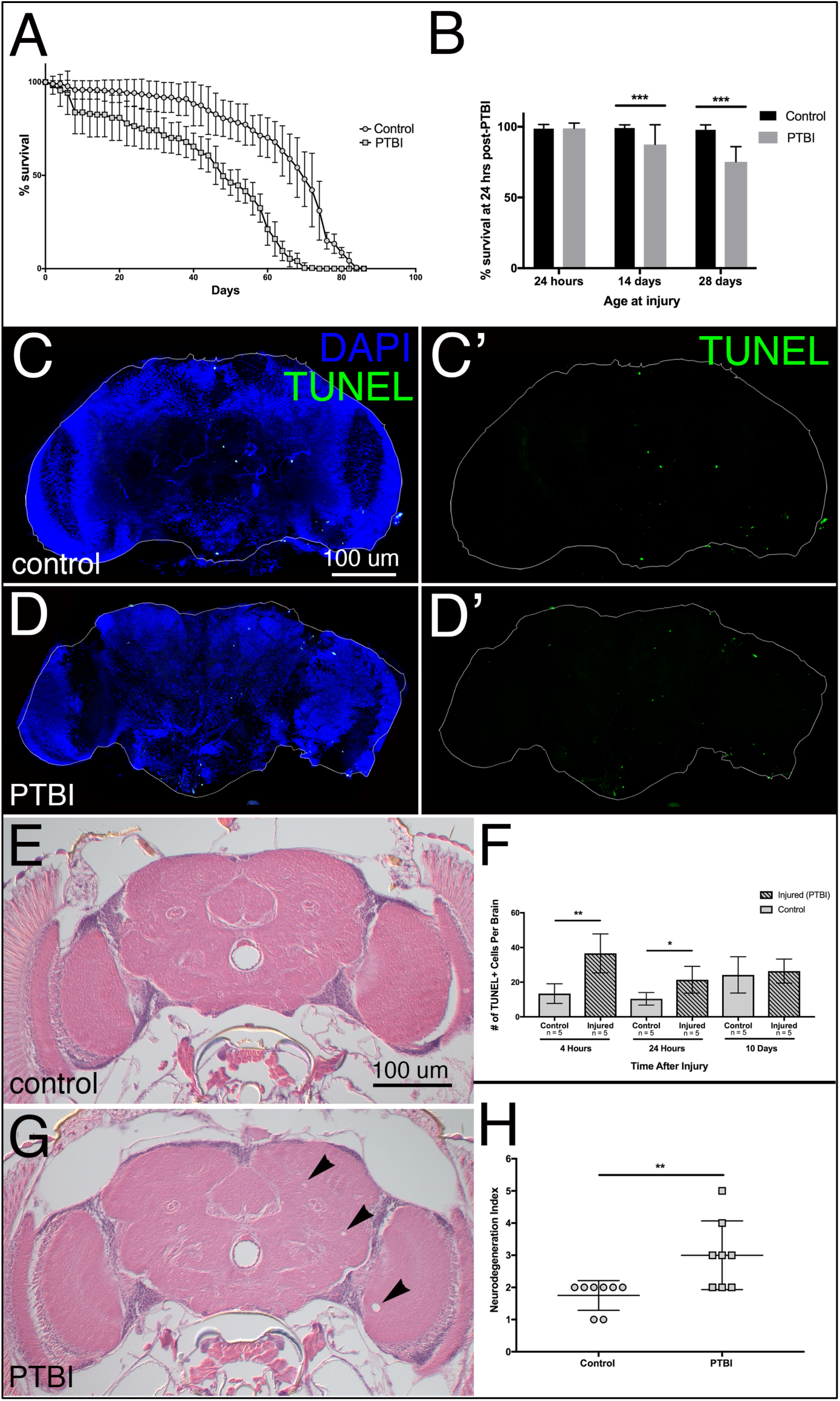
**S2A,B. PTBI decreases lifespan. A.** To assess the impact of PTBI on viability, we performed a lifespan assay with control and PTBI adult male flies. For each condition, >300 flies were assayed. Aging was carried out in vials containing 20-40 flies. Dead flies were counted and surviving flies placed on clean food every 2-3 days. Within the first 12 days, there was no significant difference in survival between control and injured flies. However, beyond 12 days, we saw a significant drop in survival of injured animals. Control males reached 50% survival at 70 days of age, while PTBI males reached 50% survival at 48 days. The maximum lifespan was 84 days for uninjured flies and 74 days for injured flies. This indicates that PTBI does impact lifespan. However, in the window of 0-12 days, there is no significant difference in survival, suggesting that death is more likely due to delayed secondary injury. Error bars reflect the standard deviation (SD). **B.** To test whether age at injury affects outcome, we compared flies injured at 0-6 hours, 14 days, and 28 days post-eclosion. Survival was assayed 24 hours later. Control flies at all ages and flies injured 0-6 hours post-eclosion, exhibited survival of 98-99% with no significant difference between age-matched control and PTBI flies (p<0.69). However, flies injured 14 days post-eclosion showed a significantly reduced survival of 88% (p<0.0001), while flies injured 28 days post-eclosion showed a further reduction to 75% survival (p<0.0001) compared to age-matched controls. These data indicate that age at the time of PTBI does affect survival. Error bars reflect the standard deviation (SD). **S2C-D’. Cell death following PTBI is increased transiently after injury.** To directly test whether PTBI causes cell death, we used terminal deoxynucleotidyl transferase dUTP nick end labeling (TUNEL). TUNEL marks the terminal stages of death, both apoptotic and necrotic, when nuclear DNA has been cleaved and degraded by DNases (Grasl-Kraupp et al. 1995). We assayed cell death using TUNEL staining (in green) in control and PTBI brains at 4 hours, 24 hours and 10 days post-PTBI. DAPI is in blue. We observe a transient increase in cell death. At 4 hours post-PTBI, there are an average of 37 TUNEL+ cells/brain (n=5 brains, 183 cells) compared to only 13 TUNEL+ cells/brain (n=5 brains, 67 cells) in uninjured controls. At 24 hours post-PTBI, there are an average of 21 TUNEL+ cells/brain (n=5 brains, 107 cells) compared to only 11 TUNEL+ cells/brain (n=5 brains, 53 cells) in uninjured controls (compare **D** and **D’** to **C** and **C’**). And at 10 days post-PTBI, there are an average of 26 TUNEL+ cells/brain (n=5 brains, 132 cells) compared to 24 TUNEL+ cells/brain (n=5 brains, 121 cells) in uninjured controls. Unpaired t tests yield p values of p<0.003, p<0.02, and p<0.7 for the 3 time points, respectively. Error bars reflect the standard deviation (SD). Thus, although the total number of dying cells increases with age in the control brains, cell death due to PTBI appears to peak shortly after the injury such that by 10 days post-injury, there was no significant difference between control and injured brains (**F**). **S2E,G,H. PTBI increases neurodegeneration.** To ask whether the observed cell death following PTBI corresponded to neurodegeneration, we used a standardized index (Cao et al. 2013) to analyze histological preparations from brains 25 days after PTBI. Controls exhibited little neurodegeneration at 25 days (**E**). In PTBI flies, we observed an increase in the number of lesions (**G**). Using the neurodegeneration index described in Cao *et al*., 2013, controls had an average neurodegeneration index score of 1.7+/-0.2, while PTBI flies had an average neurodegeneration index score of 3.0+/-0.4 (**H**). This represented a statistically significant difference (p<0.01). Error bars reflect the standard deviation (SD).

